# Polysialic Acid Presentation on Microporous Scaffolds Supports Neural Repair after Ischemic Stroke

**DOI:** 10.1101/2025.09.30.674064

**Authors:** Yunxin Ouyang, Sanchuan Che, Hannah Newman, Koravit Poysungnoen, Tatiana Segura

## Abstract

Recovery following ischemic stroke remains limited due to insufficient neural regeneration. Polysialic acid (PSA), a glycan prominently expressed during neural development, modulates neural progenitor cell (NPC) plasticity and migration, but its therapeutic potential in biomaterial-based stroke therapies remains underexplored. In this study, microporous annealed particle (MAP) scaffolds conjugated with PSA (PSA-MAP) were engineered to regulate NPC fate and promote neural tissue regeneration after stroke. PSA-MAP increased the presence of Sox2-positive progenitor cells within infarct and peri-infarct regions and elevated axonal content (NF200) in the lesion, while astrocytic and vascular coverage were not detectably changed at this early stage. In addition, 3D NPC cultures in MAP showed that tethered PSA alters NPC behavior over time, with reduced progenitor marker expression and PSA-dependent shifts in morphology, consistent with progression away from a progenitor state. Together, these data identify a glycan-forward, neuro-first repair route in which PSA-MAP enhances early neural regeneration without requiring concomitant angiogenic expansion, establishing PSA-MAP as a targeted biomaterial approach for endogenous neural repair after ischemic stroke.

## 1. Introduction

Ischemic stroke is a leading cause of long-term disability worldwide, characterized by neuronal cell death, vascular damage, and chronic inflammation, ultimately resulting in significant neurological impairment and diminished quality of life^[1]^. Despite extensive research efforts, therapeutic options remain predominantly restricted to early reperfusion therapies within narrow time windows, with minimal efficacy in stimulating regenerative processes during the subacute and chronic phases^[2]^. Therefore, therapeutic strategies aimed at promoting sustained regeneration and functional recovery in these later phases remain urgently needed^[1]^.

Neural progenitor cells (NPCs) have emerged as a critical endogenous cell type with considerable regenerative potential for stroke repair. NPCs reside primarily within specialized neurogenic niches, notably the subventricular zone (SVZ), and possess the capacity for proliferation, migration toward injury sites, and differentiation into functional neurons and glial cells^[3]^. Stroke itself induces a marked proliferation of NPCs within the SVZ, demonstrating the brain’s intrinsic regenerative response. However, these newly generated progenitors rarely migrate sufficiently far into the infarct core or its immediate vicinity^[3a, 4]^.

Rather, they predominantly remain localized within the SVZ and peri-infarct regions, greatly limiting their regenerative efficacy. Previous strategies have attempted to enhance NPC migration and differentiation by delivering growth factors, cytokines, or engineered biomaterials, often achieving improved NPC numbers in the peri-infarct area^[5]^. Nevertheless, infiltration of NPCs directly into the infarct core remains inadequate, underscoring the need for approaches that effectively guide and maintain NPCs within the stroke lesion itself.

Our prior work demonstrated that microporous annealed particle (MAP) scaffolds, composed of crosslinked hyaluronic acid (HA) microgels, could substantially improve the infiltration of endogenous NPCs into the infarct cavity, providing a supportive structural and biochemical microenvironment for cellular integration and tissue regeneration^[3b]^. Further, in our recent studies, we showed that cryo-MAP scaffolds incorporating stromal-derived factor 1 (SDF-1) nanoparticles promoted significant recruitment of SOX2-positive progenitor cells and Tuj1-positive neuronal cells into both peri-infarct and infarct regions, confirming the benefits of combining spatially controlled biomaterial presentation with biochemical signaling^[6]^. These collective findings emphasize the potential of specifically designed biomaterials to robustly enhance endogenous NPC recruitment and differentiation after stroke.

To further advance biomaterial efficacy, we have turned our attention toward polysialic acid (PSA), a glycan composed of α2,8-linked N-acetylneuraminic acid (Neu5Ac), motivated by its critical physiological roles in neural development and its potential to modulate regenerative processes within neural tissues^[7]^. PSA modifies the neural cell adhesion molecule (NCAM), altering its adhesive properties and enabling critical neural processes such as cell migration, axon guidance, and synaptic plasticity^[8]^. During brain development, PSA-NCAM expression is high, facilitating NPC migration and differentiation, axonal pathfinding, and synaptic remodeling^[7, 9]^. PSA-NCAM is found extensively on multiple cell types including neurons, glial cells, and neural progenitors, suggesting its central involvement in neural regeneration and repair processes^[7]^. PSA-NCAM’s functionality derives from the highly negatively charged PSA polymer, which, by reducing NCAM’s adhesive properties, promotes increased cellular plasticity and migration—functions vital for regenerative processes in neural tissue^[10]^. Additionally, PSA interacts directly with immune cells through inhibitory Siglec receptors, enabling immunomodulatory effects beneficial to post-injury repair^[11]^. Indeed, our recent studies demonstrated that PSA-functionalized MAP (PSA-MAP) scaffolds significantly modulated immune polarization, reducing pro-inflammatory activation of macrophages and promoting regulatory phenotypes in vitro and in vivo^[12]^. However, the direct effects of PSA-MAP on neural progenitor cells and neural cell types in the context of regenerative processes after stroke remain to be systematically investigated.

Given PSA’s extensive roles during neural development and regeneration, and its known interactions with neural and glial cell populations, we hypothesized that covalent tethering of PSA within MAP scaffolds (PSA-MAP) would further enhance NPC differentiation and neural regeneration after stroke by providing a spatially and biochemically favorable microenvironment. In this study, we evaluate whether PSA-MAP promotes neural repair after stroke, focusing on *in vivo* readouts. In a mouse photothrombotic model, we test whether PSA tethering increases progenitor presence in and around the lesion, augments axonal signal within the infarct, and modulates astroglial and vascular compartments. We then use a defined 3D NPC culture within MAP to probe how tethered PSA relates to morphology and progenitor maintenance versus loss over time within the scaffold void space. This study is designed to determine whether a stable, localized sialoglycan cue can bias the post-stroke niche toward neural repair without requiring complex protein cargo.

## 2. Results and Discussions

### 2.1. PSA-MAP Scaffolds Enhance Neural Progenitor Cell Recruitment After Stroke

To evaluate the therapeutic potential of PSA-tethered MAP scaffolds in promoting progenitor cell recruitment after stroke. We selected scaffolds containing 20 mM PSA (PSAb) to compare with HA only MAP condition (M). Photothrombotic ischemic stroke was induced on day 0, scaffolds injected into the infarct region at day 5, and animals were analyzed 10 days post-injection (day 15; **Figure 1a,b**). Quantification of Sox2-positive area demonstrated significantly higher NPC presence with PSA_b_ versus M in both the infarct (≈23% vs. 12%, **Figure 1d**) and peri-infarct (≈25% vs. 17%, **Figure 1e**) regions, indicating more robust progenitor infiltration toward and into the core at this time point.

**Figure 1.**
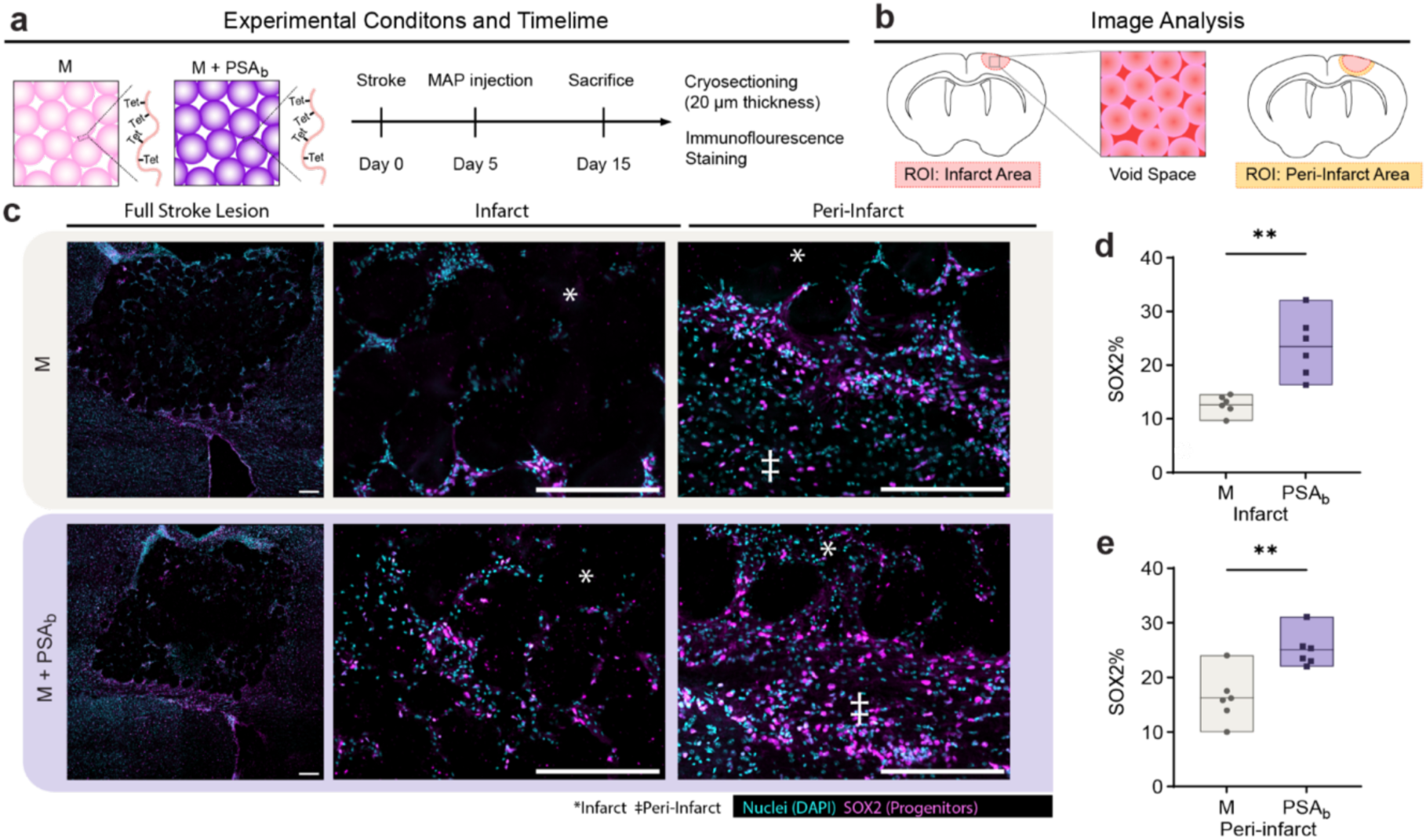
PSA-MAP scaffolds increase infiltration of SOX2-positive progenitor cells into the infarct and peri-infarct regions post-stroke. (a) Experimental timeline for in vivo stroke studies. Mice underwent photothrombotic ischemic stroke (day 0), scaffold injection into the infarct core (day 5), and were analyzed on day 15. Scaffolds were composed of microporous annealed particle (MAP) scaffolds containing either 0.5 mM RGD alone (M condition) or 0.5 mM RGD with covalently tethered 20 mM PSA (PSAb). (b) Schematic illustrating regions of interest (ROI) analyzed for immunofluorescence, including the infarct core and surrounding peri-infarct area. (c) Representative low- and high-magnification confocal images of brain sections stained for SOX2 and DAPI. (d-e) Quantification of SOX2-positive cell populations within infarct (d) and peri-infarct regions (e). Each data point is a biological replicate averaged from two coronal sections and plotted in a floating bar (n = 6). Two-tailed, unpaired Student’s t-test was performed: **p < 0.01. Scale bars represent 200 µm.

These in vivo findings align with the developmental roles of PSA in lowering NCAM-mediated adhesion to support cell motility^[8a, 10b, 13]^. In addition, our prior study showed that PSA-MAP shifted the peri-infarct compartment toward a less inflammatory, microglia-dominant state^[12]^—an environment more permissive to NPC migration and early engraftment^[14]^. Because Sox2 is not lineage-exclusive in the injured adult brain^[15]^, we interpret Sox2 here as a broad progenitor and next sought to explore the phenotypic context for this signal under more defined conditions. Thus, we conducted preliminary in vitro assays to see how PSA presentation mode and dose shape early NPC marker patterns.

### 2.2 PSA Tethering within MAP scaffolds modulate NPC morphology and differentiation *in vitro* over time

We first asked how tethered versus soluble PSA and PSA dose influence NPC markers in a simple 2D setting. NPCs were cultured five days on coverslips under the following conditions: glass, laminin, HA, HA+PSA at 1.5/15/150 µM bound, or HA with 150 µM soluble PSA. At day 5, surface-bound PSA produced dose-responsive increases in both GFAP/DAPI and Tuj1/DAPI area. HA+150 µM bound PSA significantly increased GFAP/DAPI area compared to HA, laminin, and glass (**Figure S1c**). Although the omnibus ANOVA was not significant for Tuj1/DAPI, the upward trajectory mirrored GFAP (**Figure S1d**). Presentation also mattered in this context. At 150 µM, bound PSA outperformed soluble PSA, with GFAP/DAPI significantly higher for bound (∼ 3-fold increase) and Tuj1/DAPI also higher (**Figure S1e–f**). In NPCs isolated from SVZ, Tuj1 indicates early neuronal commitment, while GFAP reflects radial-glial or progenitor states, not mature reactive astrocytes^[16]^. Thus, the coordinated rise of Tuj1 and GFAP with bound PSA suggests PSA does not prolong stemness on 2D. Instead, it nudges NPCs toward early lineage allocation in a presentation-dependent manner.

To further explore how PSA incorporation influences NPC behavior in 3D, we then conducted an *in vitro* study using a well-defined MAP encapsulation and culture protocol (**Figure S4a**). HA-HMP, either unmodified or covalently functionalized with defined concentrations of PSA and the adhesive ligand RGD, were mixed with NPCs at a cell density of 10,000 cells/µL. This mixture was annealed using a HA–Tet crosslinker, and the resulting microporous annealed particle (MAP) scaffolds were cultured for four days in vitro. Media was exchanged on day 2, and scaffolds were fixed on day 4 for subsequent immunofluorescence (IF) analysis. Five scaffold conditions were investigated: 1) no RGD and no PSA (negative control), 2) 0.5 mM RGD and no PSA (RGD-only), 3) 0.5 mM RGD with 0.625 mM PSA, 4) 0.5 mM RGD with 2.5 mM PSA, and 5) 0.5 mM RGD with 20 mM PSA. HMP size distributions showed minor differences (< 5µm) among conditions (**Figure S2c**). Fluorescence imaging confirmed a PSA-dependent increase in labeling intensity, validating controlled PSA conjugation densities (**Figure S2d**). Additionally, confocal imaging confirmed NPCs were evenly distributed within the scaffold void spaces immediately following encapsulation (day 0), indicating successful initial cell incorporation (**Figure S3**).

At day 4, NPC morphology and distribution within the scaffolds were assessed by immunostaining for the progenitor marker Sox2 and neuronal marker Tuj1, alongside phalloidin staining to visualize cytoskeletal structure (**Figure S4a–d**). In scaffolds without adhesion ligands (no RGD/no PSA), NPCs predominantly formed small, spherical aggregates, consistent with limited cell-material interactions (**Figure S4b**). In contrast, conditions containing RGD generally displayed enhanced cell spreading, indicated by decreased aggregate sphericity. We observed significant differences in aggregate sphericity among conditions, with no RGD/no PSA scaffolds having the highest median sphericity (>0.8), confirming minimal spreading due to lack of cell adhesion sites (**Figure S4e**). Interestingly, the scaffold with 2.5 mM PSA exhibited the lowest median sphericity (∼0.5), suggesting maximal cell spreading at this intermediate PSA density. Higher (20 mM PSA) and lower (0.625 mM PSA) PSA concentrations displayed intermediate spreading behaviors, potentially indicating nuanced interactions between PSA density and integrin-mediated adhesion. Analysis of Sox2-positive nuclei at day 4 revealed no statistically significant differences among scaffold conditions, with considerable variability within each group. However, an interesting observation was the overall lower Sox2 positivity (∼20%) seen at 0.625 mM PSA concentration compared to other groups (**Figure S4f**). For neuronal differentiation (Tuj1), a clear visual trend emerged, showing increased Tuj1-positive cells at higher PSA concentrations (∼18% at 0 mM PSA, rising to ∼43% at 20 mM PSA, **Figure S4g**). Although these differences did not reach statistical significance, the trend suggests PSA’s potential role in promoting neuronal differentiation at early culture stages.

To understand how these early observations evolved, scaffolds were cultured and analyzed at day 7 (**Figure 2a–d**). Characterization at this later timepoint confirmed homogeneous NPC distribution and larger aggregate size, particularly in scaffolds without adhesive ligands (no RGD/no PSA; **Figure 2b**). Aggregate sphericity quantification revealed that no RGD/no PSA scaffolds continued forming large spherical aggregates with high sphericity (median ∼0.83), reflecting sustained minimal interaction with scaffold materials. Among RGD-containing scaffolds, the highest PSA concentration (20 mM) resulted in higher aggregate sphericity (median ∼0.72), compared to lower PSA concentrations and RGD-only scaffolds (**Figure 2d**). This observation suggests possible interference by high PSA densities with integrin-mediated cell spreading, leading to partially reduced cellular elongation. Importantly, significant PSA-dependent differences in Sox2 expression emerged at day 7. The RGD-only condition retained high Sox2 expression (∼65%), suggesting sustained progenitor identity, whereas PSA-containing scaffolds showed marked reductions (∼14% at 0.625- and 2.5-mM PSA, and <10% at 20 mM PSA). Statistical significance was observed when comparing PSA groups to both the RGD-only and no RGD/no PSA conditions, strongly indicating that tethered PSA effectively reduced progenitor marker expression, potentially indicating enhanced differentiation (**Figure 2f**). Analysis of Tuj1 expression at day 7 further supported the trend observed at day 4, showing a clear visual increase in neuronal differentiation markers at higher PSA concentrations, with the highest PSA group (20 mM) exhibiting ∼40% Tuj1-positive cells. Lower PSA concentrations and RGD-only groups showed lower Tuj1 expression (∼10%) (**Figure 2g**). While statistical comparisons did not confirm definitive differences, the clear directional trend supports PSA’s potential involvement in regulating NPC differentiation, consistent with PSA’s established developmental involvement^[8d, 17]^.

**Figure 2.**
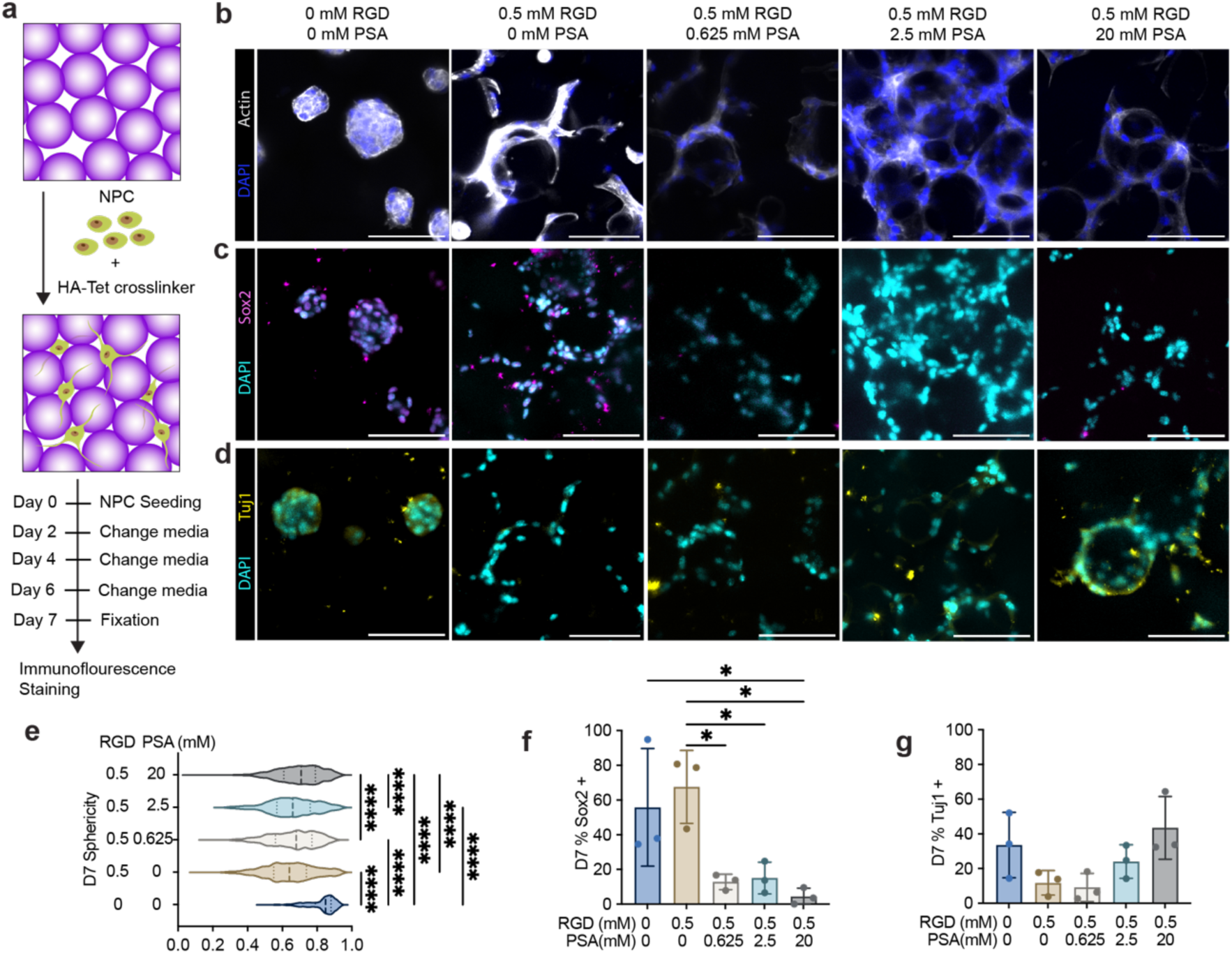
Effects of scaffold-tethered PSA on NPC morphology and progenitor marker expression at day 7. a) Schematic of experimental setup illustrating NPC encapsulation within MAP scaffolds for a 7-day culture period. b–d) Representative confocal images of NPCs in scaffolds stained with DAPI and actin (b), the progenitor marker SOX2 (c), or early neuronal differentiation marker Tuj1 (d). Scale bars: 100 µm. e) Violin plots quantifying cell aggregate sphericity across scaffold conditions. Each data point represents an individual aggregate (>200 aggregates per condition, pooled from three biological replicates). f,g) Quantification of the percentage of SOX2-positive (f) and Tuj1-positive (g) nuclei. Data represent mean ± SEM from three biological replicates, where each replicate corresponds to NPCs thawed from separate vials, scaffolds prepared independently, and experiments conducted on separate days (n = 3 scaffolds per condition). Statistical significance determined by one-way ANOVA with Tukey’s post-hoc test (*p < 0.05, **p < 0.01, ***p < 0.001, ****p < 0.0001).

Collectively, these findings reveal a dynamic, PSA-dependent shift in NPC behavior within MAP scaffolds over the seven-day culture period. In vitro, PSA preserves a progenitor-like state only transiently—Sox2 is maintained early and then declines by ∼day 7 as differentiation initiates. These in vitro findings provide insights to our vivo observations–the elevated Sox2 signal we observe in vivo at day 15 most likely reflects continued recruitment and short-term persistence of NPCs within PSA-MAP, rather than PSA ‘locking’ early cell entrants in a stem-like state.

### 2.3. PSA-MAP Scaffolds Promote Axonal Regeneration Without Altering Astrocytic Responses

PSA-MAP’s pro-NPC effects led us to ask whether these cues also translate into structural neural repair at day 15 post-stroke (**Figure 3a,b**). GFAP-positive area was similar between groups in the peri-infarct, whereas within the infarct the PSA-MAP group exhibited a slightly higher mean GFAP-positive area with no statistical significance (**Figure 3c–e**). In contrast, NF200, an axonal marker, increased significantly within the infarct core with PSA-MAP (≈15% vs. ≈7% with MAP-only) and showed a modest, non-significant elevation in the peri-infarct (**Figure 3f–h**). Collectively, these data indicate that PSA-MAP enhances axonal entry/sprouting into the lesion cavity without a significant change in astrocyte coverage at this time point.

**Figure 3.**
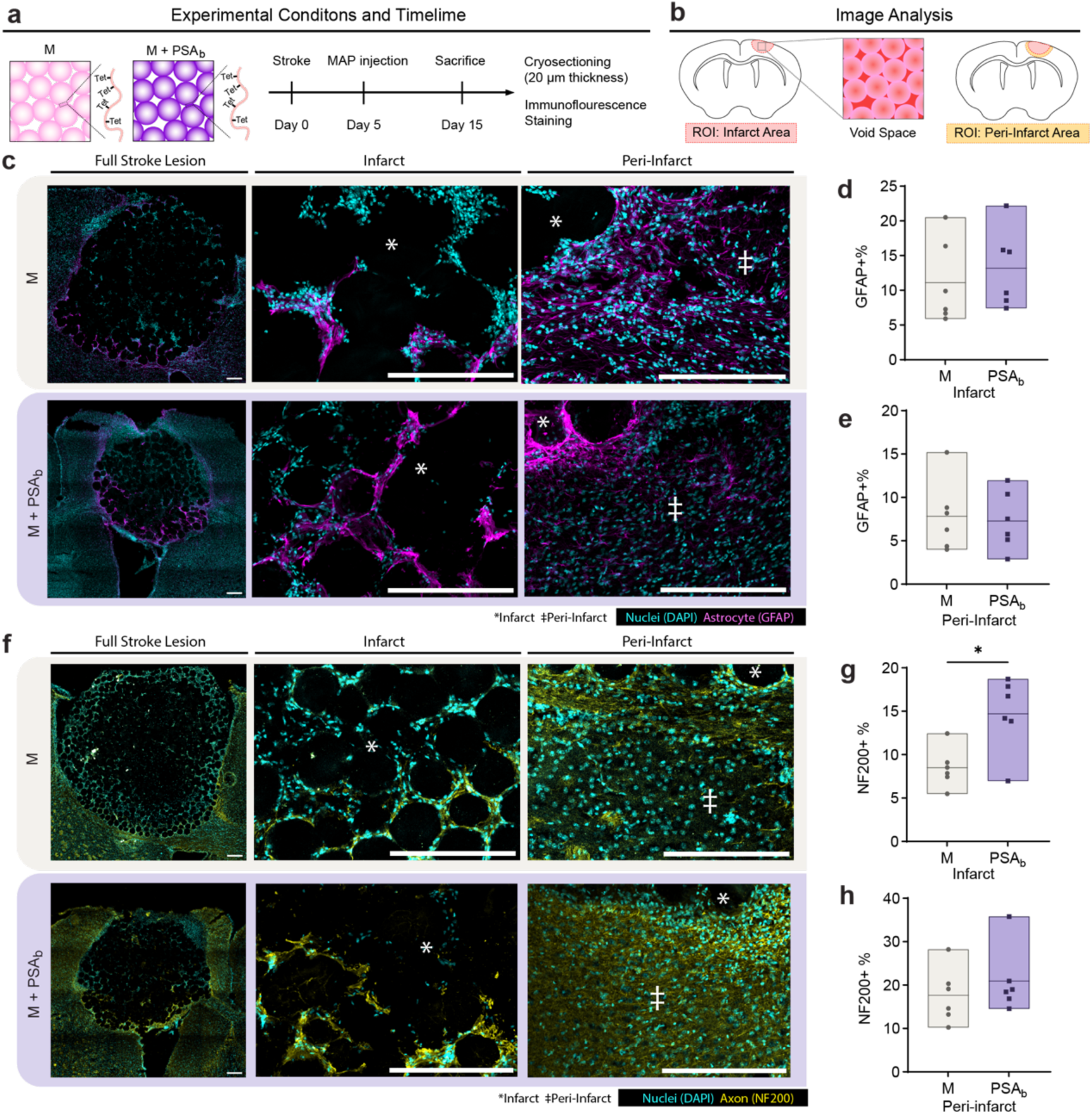
PSA-MAP scaffolds enhance axonal regeneration but do not significantly alter astrocytic infiltration post-stroke. (a,b) Experimental timeline and image analysis ROIs identical to Figure 3. (c) Representative confocal images of GFAP staining in infarct and peri-infarct regions. (d,e) Quantification of GFAP-positive area in infarct (d) and peri-infarct (e) regions. (f) Representative confocal images of NF200 staining indicating axonal infiltration. Scale bars: 100 µm. (g,h) Quantification of NF200-positive area in infarct (g) and peri-infarct (h) regions. Each data point is a biological replicate averaged from two coronal sections and plotted in a floating bar (n = 6). Two-tailed, unpaired Student’s t-test was performed: **p < 0.01. Scale bars represent 200 µm.

The NF200 increase aligns with PSA’s biophysical role in lowering NCAM-mediated neuronal adhesion and promoting plasticity: tethered PSA can reduce intercellular adhesion barriers and bias the local niche toward axon extension and motility^[9b, 18]^. Reactive astrocytes have dual roles after injury: while scar architecture can restrict cellular movement[19], astrocytes also contribute critically to tissue stabilization and vascular function (e.g., supporting BBB integrity and water/ion homeostasis via endfeet specializations such as AQP4)[20]. Thus, the slightly higher GFAP signal within the infarct in the PSA-MAP group may reflect local differences in astrocyte organization or maturation that are compatible with axonal ingress.

### 2.4 PSA-MAP Scaffolds Show Limited Effects on Early-stage Vascular Remodeling After Stroke

Given the neural benefits at day 15, we next asked whether these neural gains required concurrent angiogenesis (**Figure 4a–b**). In both infarct and peri-infarct, CD31 (pan-endothelium) showed small yet non-significant increases with PSA-MAP (**Figure 4c–e**), whereas glut1, a glucose transporter enriched in brain microvascular endothelium, did not differ between groups (**Figure 4f–h**). In other words, at this early subacute time point we detect clear neural enhancements without a measurable change in endothelial area or BBB-associated phenotype by area metrics.

**Figure 4.**
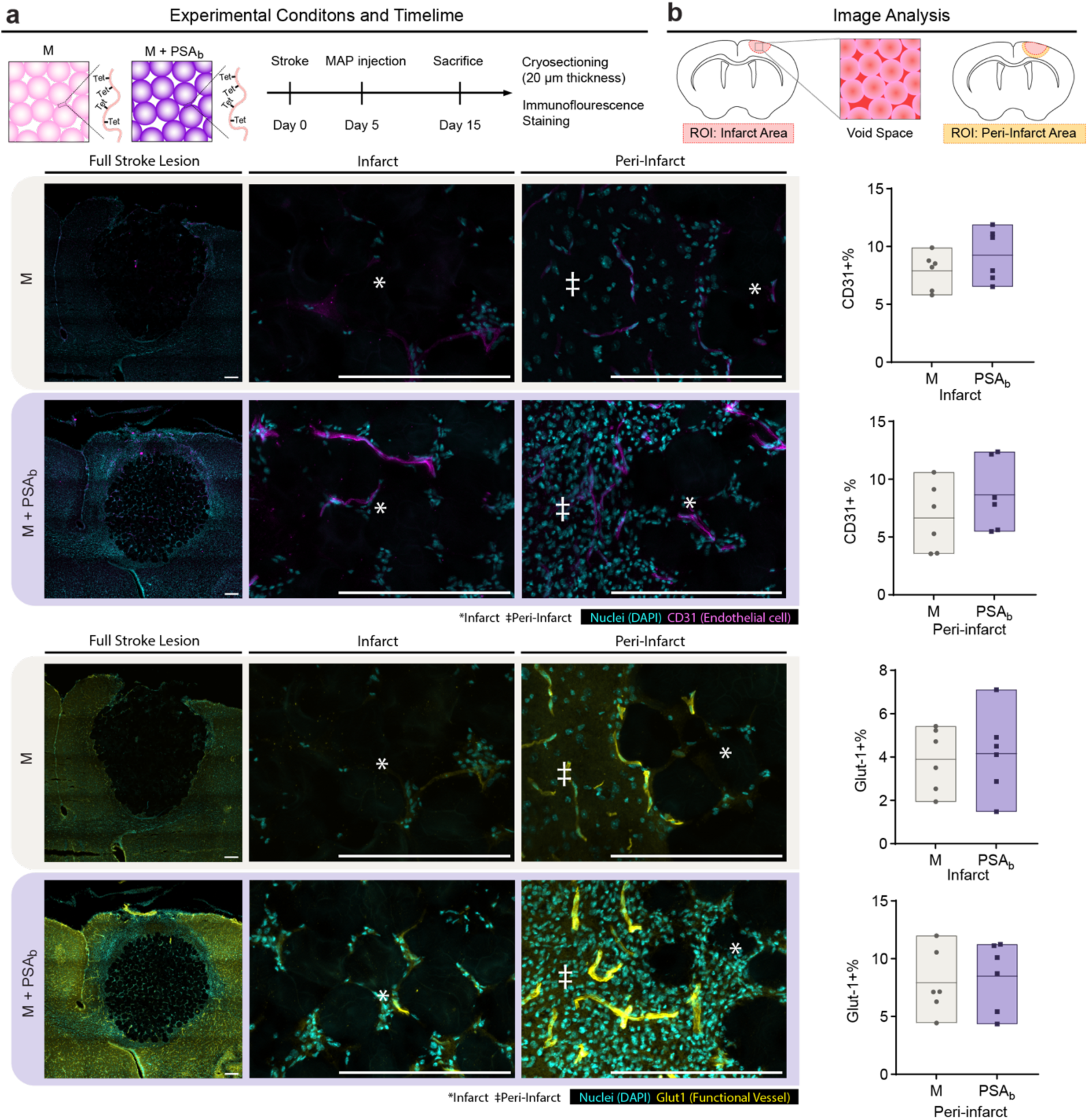
Early neural gains with PSA-MAP occur without a detectable early angiogenic shift at day 15. (a,b) Experimental timeline and image analysis ROIs consistent with Figures 3 and 4. (c) Representative images of CD31 staining for general endothelial cell visualization. Scale bar: 100 µm. (d,e) Quantification of CD31-positive area in infarct (d) and peri-infarct (e) regions. (f) Representative images of GLUT1 staining, indicative of mature vessels. Scale bar: 100 µm. (g,h) Quantification of GLUT1-positive area in infarct (g) and peri-infarct (h) regions. Each data point is a biological replicate averaged from two coronal sections and plotted in a floating bar (n = 6). Two-tailed, unpaired Student’s t-test was performed: **p < 0.01. Scale bars represent 200 µm.

This pattern differs from a prior biomaterial system from our lab in which a nanoporous HA hydrogel containing CLUVENA drove robust angiogenesis that was tightly coupled to axonal ingrowth and functional recovery; pharmacological blockade of angiogenesis with endostatin in that system abrogated both axonal gains and behavioral improvement, indicating an angiogenesis-dependent route to neural repair[21]. Here, by comparison, PSA-MAP yields significant increases in Sox2 and NF200 in the absence of a detectable early increase in CD31 or Glut1 area, suggesting that neural plasticity can be initiated in this scaffold without an obligate, preceding expansion of angiogenic area at day 15. Mechanistically, PSA-MAP primarily acts through neural contacts—reducing NCAM-mediated adhesion and facilitating progenitor motility[17b] and axon extension[9b]— while also softening inflammatory tone, thereby unmasking neural programs that do not strictly require an early angiogenic burst. The VEGF-nanoparticle hydrogel, in contrast, was designed to amplify endothelial growth directly; in that context, vessels provided both structural and trophic support to axons, and blocking angiogenesis removed that support.

Taken together, the data indicate that PSA-MAP can deliver early, measurable neural gains without a detectable early angiogenic shift. This neuro-first sequence is distinct from the angiogenesis-first dependency observed in the CLUVENA hydrogel and highlights that different material cues can prioritize different entries into the neurovascular repair loop.

To complement area-based readouts, we additionally quantified infiltration distance (the depth that positive signal extended from the cavity boundary into the infarct) for Sox2, NF200, GFAP, and CD31 (Glut1 was not analyzed due to low infarct signal). Across all four markers, the PSA-MAP means trended higher than MAP-only, but none reached statistical significance in this cohort (**Figures S8-S11**). These trend-level increases are directionally consistent with the significant gains observed in area quantification, yet the distance metric proved more variable. The greater variance likely reflects sensitivity of the distance metric to lesion fill fraction, microgel packing, and RGD presentation—parameters that can be further standardized in future experiments.

## 3. Conclusion

In this study, we evaluated PSA-MAP scaffolds, previously shown to modulate immune responses[12], for their ability to influence neural regeneration and neurovascular remodeling after ischemic stroke. Our motivation stemmed from the critical clinical need to enhance endogenous regeneration during the subacute stroke phase, coupled with PSA’s well-established developmental and regenerative roles within neural tissues.

Using a mouse photothrombotic stroke model, we demonstrated significant beneficial neural outcomes from PSA-MAP scaffolds. PSA-tethered scaffolds markedly increased infiltration of SOX2-positive progenitor cells into infarct and peri-infarct regions, directly supporting our hypothesis that PSA conjugation enhances endogenous progenitor recruitment. This enhanced NPC infiltration aligns closely with PSA’s known developmental roles in facilitating progenitor cell migration and supports the scaffold’s regenerative potential^[8e]^. Furthermore, *in vitro* findings provided additional insights on how PSA tethering influenced NPC phenotype. In the 3D MAP scaffold, PSA initially maintained progenitor identity but over time promoted early differentiation. *In vivo*, PSA-MAP scaffolds significantly promoted axonal regeneration, as evidenced by increased NF200-positive areas within infarct sites. These neural-specific regenerative effects underscore PSA’s strong neural-regulatory capacity^[7]^. Interesting, these neural gains were observed alongside largely unchanged GFAP area and no measurable increase in CD31 or GLUT1 area, indicating that early benefits did not depend on a detectable, concomitant expansion of endothelial area by our metrics. Together, these data support an early, neuro-first repair sequence in PSA-MAP: increased progenitor ingress and axonal signal can be achieved without a required, early angiogenic shift.

The findings that PSA scaffolds robustly increased NPC infiltration and axonal regeneration yet exhibited limited effects on early vascular remodeling and astrocytic responses, also offer considerations for future scaffold design. Combining PSA conjugation with additional design factors known to robustly stimulate angiogenesis such as the cryoMAP, may represent an effective approach to simultaneously stimulate both neural and vascular regeneration[6].

Manipulating scaffold porosity or void fraction could further optimize vascular ingrowth and cell migration, improving overall tissue integration. Indeed, a combined PSA–CryoMAP scaffold approach, leveraging both controlled PSA presentation for neural modulation and controlled void fraction with angiogenic nanoparticle delivery, represents an attractive strategy for promoting simultaneous neurogenesis and angiogenesis in stroke repair. Long-term evaluations of PSA-MAP scaffolds remain an important next step. The current study focused on relatively early regenerative responses following scaffold implantation.

Investigating longer-term tissue repair, maturation of neurons, glial remodeling, and sustained vascular maturation will be critical for understanding therapeutic outcomes at chronic stages[22]. Additionally, future functional outcome studies—including motor, sensory, and cognitive assessments—would further elucidate whether observed cellular and morphological improvements translate into measurable functional recovery[23]. The integration of behavioral testing with histological and molecular analyses could greatly enhance the translational relevance of our findings[22a].

## 4. Experimental Section/Methods

### Modification of Hyaluronic Acid with Norbornene

HA–norbornene (HA–NB) was synthesized by activating the carboxyl groups of hyaluronic acid (79 kDa, Contipro) using 4-(4,6-dimethoxy-1,3,5-triazin-2-yl)-4-methylmorpholinium chloride (DMTMM; TCI America). HA and DMTMM were each dissolved in 200 mM MES buffer (pH 5.5) and combined at a molar ratio of 1:4 (HA repeat unit:DMTMM). After stirring for 10 min at room temperature, 5-norbornene-2-methanamine (NMA; TCI America) was added dropwise (2 molar equivalents per HA repeat unit). The mixture was left to react overnight at room temperature. The reaction product was precipitated in cold ethanol (1 L,4 °C), and the resulting solids were collected. To remove unreacted small molecules, the precipitate was dissolved in 2 M brine solution and dialyzed sequentially against 2 M brine, 1 M brine, and finally DI water (MWCO 12–14 kDa). The dialysis cycle was repeated three times, followed by a 24 h dialysis against DI water. The purified solution was lyophilized to yield the final HA–NB polymer. Successful functionalization was verified by ^1^H NMR (D₂O), with characteristic norbornene vinyl proton resonances at δ 6.33 and δ 6.02 (endo) and δ 6.26 and δ 6.23 (exo). Comparison with the HA methyl resonance at δ 2.05 ppm indicated ∼31– 45% substitution of HA repeat units (Figure S4). The resulting HA–NB served as the base macromer for HMP fabrication and subsequent MAP scaffold assembly.

### Modification of Polysialic Acid Reducing End with Thiol and Tetrazine

PSA functionalization was carried out through a reductive amination approach adapted from prior methods[24] and described in detail in our companion immune-focused study^[25]^. Briefly, polysialic acid (MW 24–38 kDa; Santa Cruz Biotechnology) was first purified to remove endotoxins and then derivatized at its reducing end with cystamine, followed by reduction and thiol liberation to generate PSA–SH. To enable site-specific and orthogonal conjugation, PSA–SH was converted to PSA–tetrazine (PSA–Tet) via thiol–maleimide Michael addition. The resulting product was purified by sequential dialysis and lyophilization. Successful functionalization was confirmed by NMR, which showed the characteristic tetrazine aromatic proton signals at δ 8.35 and δ 7.20 ppm in D₂O (Figure S5). This strategy ensured stable PSA presentation through bio-orthogonal chemistry, allowing subsequent conjugation to HA–NB microparticles while preserving a physiologically relevant glycan orientation.

### HMP Production and Purification

HA–NB hydrogel microparticles (HMPs) were generated using a planar flow-focusing microfluidic device to produce uniform droplets. The aqueous precursor consisted of hyaluronic acid–norbornene (HA–NB), an MMP-sensitive dithiol peptide linker (Ac-GCRDGPQGIWGQDRCG-NH2), lithium phenyl-2,4,6-trimethylbenzoylphosphinate (LAP) photoinitiator, and, where indicated, RGD adhesive peptide. Droplets were pinched off by a continuous oil phase containing surfactant and subsequently exposed to UV light (365 nm) to initiate thiol–norbornene crosslinking, yielding stable HMPs. Following polymerization, emulsions were broken and particles were repeatedly washed with buffer to remove fluorinated oil and surfactant. Centrifugation steps were performed until no residual oil was visible. Particle size distributions (≈70–100 μm) were confirmed by microscopy (Figure S2a– b). Endotoxin testing confirmed final preparations were consistently below 0.2 EU mL⁻¹. PSA functionalization was achieved by reacting HMPs with tetrazine-modified PSA, enabling conjugation to unreacted norbornene groups on the particle surface. Uniform PSA presentation was verified using fluorescently labeled PSA (Figure S2c–d). The full chemical formulation and additional purification details are provided in our companion immune-focused paper^[25]^.

### HMP Post-Fabrication Modification with PSA-SH-Tet

For *in vitro* cell culture and *in vivo* injection, HA–NB HMPs were modified incubated in 0.22µm sterile filtered 3 × 10^−1^ M HEPES pH 7.4 containing PSA-SH-Tet at final Neu5Ac monomer concentrations of 0 M, 1.25 x 10^-3^ M, 2.5 x 10^-3^ M, or 2 x 10^-2^ M at 37 °C for 4 h with agitation every 1 h. The suspended HMPs were then pelleted by centrifuging at 14 000 rcf for 15 min. The HMPs were washed three times in a laminar flow tissue-culture hood with sterile filtered 3 × 10^−1^ M HEPES, one time wash with 70% EtOH for 8 min, three times again with sterile filtered 3 × 10^−1^ M HEPES, and recovered with the same centrifugation conditions as above. The recovered HMPs were then exposed to UV sterilization for 2 hours before in *vivo* and *in vitro* applications. PSA conjugation efficiency quantification was performed as previously reported^[12]^. Briefly, the HMPs were incubated in 3 × 10^−1^ M HEPES pH 7.4 containing AF555–PSA-Tet (synthesized in-house via non-reducing end oxidation and aniline catalyzed oxime ligation, as previously reported) at initial Neu5Ac monomer concentrations of 0 M, 6.25 x 10^-4^ M, 2.5 x 10^-3^ M, or 2 x 10^-2^ M at 37 °C for 4 h with agitation every 1 h. The suspended HMPs were then pelleted by centrifuging at 14 000 rcf for 15 min. The HMPs were washed three times with 3 × 10^−1^ M HEPES and recovered with the same centrifugation conditions. Modified HMPs were imaged on a Nikon Ti Eclipse with a 20× air objective and quantified via Cellpose Cyto3 model version 3.0.10, as previously reported^[12]^.

### Modification of Hyaluronic Acid with Tetrazine

HA–tetrazine (HA–Tet) was synthesized following a carbodiimide-mediated coupling strategy adapted from previous reports [25] and described in full detail in our companion immune-focused study[12]. In brief, hyaluronic acid (79 kDa) was activated with DMTMM and reacted with tetrazine-amine to yield polymers bearing tetrazine groups on ∼4–6% of repeat units. Modified polymers were purified by dialysis (MWCO 12–14 kDa) and lyophilized. The degree of functionalization was assessed by ^1^H NMR, with diagnostic tetrazine aromatic proton signals appearing at δ 8.5 and δ 7.7 ppm, relative to the HA methyl resonance at δ 2.05 ppm (Figure S6).

### NPC Isolation

P12 C57BL/6J pups were anesthetized, and their brains were extracted. Subventricular zone (SVZ) wholemount from the brains were dissected and placed in DMEM/F12 (DF) media plus 100 units/ml penicillin, 100 μg/ml streptomycin, and 250 ng/ml amphotericin B. The dissected tissues were incubated in 0.005% trypsin (15 minutes, 37°C, and pH 7.3) and placed overnight in uncoated T75 plastic tissue-culture dishes in N5 medium; containing 35 μg/ml bovine pituitary extract, abx, 5% FCS (HyClone), N2 supplements, 40 ng/ml EGF, and basic FGF. Unattached cells were collected and subsequently plated again onto a 6 well TC treated plate. These cells were proliferated to confluency in N5 media. Fresh EGF and FGF (20 ng each) were added every other day. The cells were proliferated to 85–95% confluency, until neurospheres formed on the plate, at which point they were trypsinized, collected and frozen until further use. These cells were marked as passage 2 (P2). Generally, cells were passaged two or three times by using 0.005% trypsin (Corning) and N5 media, with 20 ng/ml EGF/FGF supplementation every other day.

### NPC Encapsulation in MAP

NPC were detached from 6-well plates 0.005% trypsin (Corning). The cell pellet was prepared at a final concentration of 10 000 cells µL^-1^ of MAP scaffolds. Subsequently, the HMP mixture (50 µL) containing HA-Tet (HA-Tet/HA-NB ratio = 0.35) was added to the cell pellet and thoroughly mixed on ice before pipetting onto a MAP-seeding chamber (MSC) to create a hemispherical dome and incubated at 37 °C for 45 min before adding 100 µL of N5 media. MSC’s negative mold was produced on a Form 2 stereolithography 3-D printer (Formlabs, Inc.; Duke SMIF). Each well features a 4 mm-diameter × 1 mm-high culture cavity capped by a 5 mm-diameter × 3 mm-high media reservoir that holds ≥ 50 µL. The mold was cast in PDMS, cured, and the finished wells were plasma-bonded to glass coverslips. The N5 media was replenished every two days by gently aspirating 60 µL of old N5 media and adding 80 µL of fresh N5 media to each well.

### NPC culture on glass coverslip

12mm glass coverslips (neuVitro) were coated in three sequential layers to generate five surface chemistries: (1) laminin control (PLO → laminin), (2) RGD only (PLO → HA-NB → RGD), and (3–5) PSA + RGD at three PSA doses (PLO → HA-NB → PSA + RGD; final PSA 1.5, 15, or 150 µM with RGD fixed at 300 µM). All steps were performed sterilely on Parafilm sheets in a biosafety cabinet, with PBS washes (3 × 5 min) between layers. 80–100 µL of each coating solution was used per coverslip. Coverslips were first coated with 500 µg mL⁻¹ poly-L-ornithine prepared from a 2 mg mL⁻¹ stock in sterile PBS. Drops were first applied to a Parafilm sheet, coverslips (cell side down) were then placed onto the drops, and incubated overnight at 4 °C, followed by PBS washes. For conditions 2–5, a HA layer was applied using HA-NB dissolved fresh at 14.64 mg mL⁻¹ in 0.3 M HEPES and sterile-filtered (0.22 µm). 80–100 µL of HA-NB solution applied in the same way as PDL and incubated overnight at 4 °C or RT for 2–4 h. The HA layer was omitted for the laminin control. PSA and RGD Ligands were then presented in the third layer. RGD (Ac-RGDSPGERCG-NH2, Genscript) was prepared as a 50 mM stock in H₂O. For the RGD-only surface, an RGD coating solution was prepared at 300 µM RGD with 9.9mM LAP in 0.3M HEPES, sterile-filtered, applied to coverslips (80–100 µL) on Parafilm, then UV-exposed for 1 min to induce SH-NB reaction. For PSA + RGD surfaces, PSA-SH was prepared at a stock concentration of 300 µM in 0.3 M HEPES and combined with 300 µM RGD and 9.9.mM LAP in 0.3M HEPES to yield final PSA concentrations of 1.5, 15, or 150 µM. Each solution was sterile-filtered in the tissue-culture hood, applied to coverslips (80–100 µL), and the surface was UV-exposed for 1 min to induce SH-NB reaction. Prior to NPC seeding, all coverslips were transferred to 24-well plates (coating side up) and rinsed 3 × 5 min with PBS, then equilibrated in NPC medium. For cell seeding, NPC were detached from 6-well plates 0.005% trypsin (Corning). Cell suspension containing 0.076 k cells µL⁻¹ were prepared, and 500 µL of cell suspension were added to each well. Media change was performed every two days.

### Immunostaining and imaging of NPC in MAP

NPC cultures were gently aspirated of media and fixed at day 4 or day 7 with 4% PFA for 30 min at room temperature and then washed with 1x PBS 3x for 15 min per wash before immunostaining. Cultures were first incubated with blocking solution containing 1x TBS, 0.3% Triton-X, and 10% normal donkey serum (NDS) for 2 hr in room temperature, then incubated with primary antibodies at the appropriate dilutions in blocking solution for overnight at 4°C. After 3x 5-minute washes in 1x PBS, with 0.3% Triton-X, samples were incubated with secondary antibodies at the appropriate dilution in clocking solution for 2 hours at room temperature. The samples were then counterstained with nuclear marker DAPI (1:500, Invitrogen) for 15 minutes at room temperature and then washed and kept in 1x PBS until imaged. *Primary antibodies*: 1:100 rabbit anti-SOX2 (Synaptic Systems, 347 003) for maintained stem cell marker, 1:100 rabbit anti-beta III Tubulin (Abcam, ab18207) for neuronal cell marker differentiation. *Secondary antibodies*: 1:500 Alexa Fluor 555 (Rb, Invitrogen, A-31572), 1:500 iFlour 647 Phalloidin (Cayman Chemical Company, 20555). All fluorescence imaging was performed using a Nikon Ti Eclipse microscope equipped with a C2 laser with a 20× air objective at a 4 µm step-size.

### IMARIS analysis of NPC in MAP

IMARIS software was used for processing and analyzing the 20x confocal images. To correct for background signal, the IMARIS masking function was first used to eliminate all Tuj1 signal that did not overlap with Phalloidin signal and all Sox2 signal that did not overlap with DAPI signal. Subsequently, 3D volumes were rendered around Tuj1, Sox2, Phalloidin, and DAPI signal. To quantify the percentage of microtubules that tested positive for Tuj1, the total volume of the 3D volumes around all Tuj1 signal was divided by the total volume of the 3D volumes around all Phalloidin signal. Similarly, to quantify the percentage of the nuclei that tested positive for Sox2, the total volume of the 3D volumes around all Sox2 signal was divided by the total volume of the 3D volumes around all DAPI signal. To measure the sphericity of the nuclei, the built-in IMARIS sphericity function was used, defining sphericity according to Wadell as the ratio of the surface area of a sphere to the surface area of a particle with the same volume as the sphere. The software provided sphericity values for each of the 3D surfaces rendered around DAPI signal. The number of nuclei in each NPC cluster was quantified using the Split Spots Into Surface Objects IMARIS Xtension. Using this Xtension, the Surface component was defined as the 3D volumes created around Phalloidin signal and the Spots were defined as the 3D volumes created around DAPI signal. Further, the option to model Point Spread Function (PSF)-elongation along the Z-axis was selected to correct for image distortion in the Z-direction resulting from using confocal microscopy without deconvolution. The spots were then separated into clusters based on which Phalloidin surface they overlap with.

### Cellpose Analysis of NPC in MAP

To generate images suitable for segmentation, maximum intensity projections (MIPs) were constructed in ImageJ by merging five consecutive z-stacks, thereby enhancing the resonant resolution of cellular features. Nuclei were segmented using the re-trained Cellpose Cyto3 model (v3.0.10) (https://doi.org/10.1038/s41592-020-01018-x), optimized on a curated dataset of over 50 manually annotated nuclear masks for improved performance in MAP-embedded cell types. Segmentation was performed using a flow threshold of 0.8 and a cell probability threshold of −1.0. Post-segmentation filtering excluded non-nuclear artifacts based on both area and DAPI intensity, using a z-score threshold of 1.0. SOX2 and Phalloidin fluorescence expression levels were quantified by computing the mean fluorescence intensity within each segmented nucleus, with SOX2 serving as the primary marker for downstream analysis. Expression values were Z-normalized across all nuclei. SOX2-positive regions of interest (ROIs) were identified via threshold-based classification on the normalized intensity data. All final overlays and ROI visualizations were rendered using NAPARI (https://napari.org/stable/).

### Photothrombotic Stroke

Photothrombotic stroke induction was performed in adult male C57BL/6 mice (8–12 weeks, Jackson Laboratories) under isoflurane anesthesia, following protocols approved by the Duke IACUC. Briefly, after Rose Bengal injection and laser illumination through the intact skull, focal cortical infarcts were generated as previously described in our companion study [12]. In the present work, all animals received scaffold injections on day 5 post-stroke, and brains were collected on day 15 for histological analysis of neural endpoints.

### Stereotactic MAP Injection

On day 5 post-stroke, a burr hole was drilled at the cortical site corresponding to the laser illumination target, and MAP scaffolds (with or without PSA tethering) were injected stereotactically into the infarct cavity. The incision was sealed using Vetbond, and mice were monitored during recovery. A complete description of stereotactic injection parameters and scaffold preparation has been published in our companion immune-focused study [12].

### Tissue Processing for Immunostaining

At the terminal endpoint (day 15), mice were anesthetized with isoflurane and perfused with at least 10 mL of ice cold 1x PBS followed by ice cold 4% (w/v) paraformaldehyde (PFA, Electron Microscopy Sciences). Following perfusion, brains were extracted and placed in a 4% PFA solution for 4 h at 4 °C. Brains were then washed three times with 1X PBS, after which they were placed into a 30% sucrose solution prepared in 1X PBS at 4 °C. Brains were cryosectioned into 30 µm slices, collected onto glass slides, and preserved at −80 °C until immunostaining.

### Immunostaining and imaging of brain sections

Immunofluorescence staining was performed as previously described in our companion study [12], with minor adjustments for neural markers. Briefly, sections were blocked in 1× TBS containing 0.3% Triton-X and 10% normal donkey serum, incubated with primary antibodies overnight at 4 °C, and exposed to fluorescent secondary antibodies and DAPI for 2 h at room temperature. After thorough washing, slides were dehydrated through graded ethanol, cleared in xylene, and mounted in DPX. *Primary Antibodies*: 1:200 SOX2 (Synaptic Systems, 347 003), 1:250 GFAP (Fisher Scientific, 13-030-0), 1:200 NF250 (Sigma-Aldrich, N4142-25UL), 1:200 Glut-1 (EMD Millipore Corp, 07-1401), 1:100 CD31 (BD Pharmingen, cat.553370). *Secondary Antibodies*: 1:500 Alexa Fluor 488 (Guinea Pig, Invitrogen, A-21450), 1:500 Alexa Fluor 555 (Rb, Invitrogen, A-31572), 1:500 Alexa Fluor 647 (Rb, Invitrogen, A-31573). All fluorescence imaging of brain sections was performed using a Nikon Ti Eclipse microscope equipped with a C2 laser with a 20× air objective at a 4 µm step-size.

### Image Analysis

Fluorescent imaging was performed using a Nikon Ti Eclipse scanning confocal microscope equipped with a C2 laser system and either 4× or 20× air objectives. Image acquisition and analysis were conducted using Fiji/ImageJ. Images are presented as maximum intensity projections (MIPs). For quantification of positive signal areas, images were first despeckled and thresholded, followed by measurement within defined regions of interest (ROIs). Signal area normalization was achieved by dividing the positively stained area by the void space, determined through thresholding of Gaussian-blurred DAPI images (σ = 2 for nuclear markers; σ = 8 for cytoplasmic markers). For infiltration measurements, linear measurements per tissue section were obtained at ≈150 um intervals. Two tissue sections per animal were analyzed for quantification.

### Statistical Analysis

NPC *in vitro* experiments were repeated to contain three biological replicates, and each biological replicate contained HMPs generated from a different microfluidics batch and NPCs thawed and passaged on different days. Statistical significance was assessed using a one-way ANOVA with Tukey post-hoc test. For in vivo immunofluorescence quantification, the sample size was determined a priori with G*Power v3.1.9.6 (two-tailed, unpaired Student’s t-test, α = 0.05, power = 0.80). Pilot data yielded an effect size of Cohen’s d = 1.77, indicating that 6 animals per group were required. For each animal, values from two brain sections were averaged, and this mean served as a single biological replicate. Results are plotted as floating-bar graphs (min–max) with individual animal means overlaid. Homogeneity of variance was confirmed (Levene’s test, p > 0.05), and group differences were evaluated with a two-tailed, unpaired Student’s t-test using the pooled-variance estimate (α = 0.05).

## Acknowledgements

The authors would like to thank Dr. Katrina Wilson and Dr. Moawiah M. Naffaa for teaching and assisting with NPC isolation. The authors would like to thank Dr. Andrea Jones for kindly helping with outlining and editing the manuscript. Lastly, we would like to acknowledge our funding from the National Institutes of Health through R01NS079691 (TS) and R01NS112940 (TS).

## Conflicts of Interest

Y.O. and T.S. are co-inventors on a provisional patent application (U.S. Provisional Patent Application No. 63/784,550), filed by Duke University, entitled ‘GLYCAN ACTIVE MICROPOROUS ANNEALED PARTICLE GEL SYSTEM AND METHOD THEREOF,’ which relates to the polysialic acid-functionalized MAP technology described in this manuscript. The other authors declare no competing interests.

Received: ((will be filled in by the editorial staff))

Revised: ((will be filled in by the editorial staff))

Published online: ((will be filled in by the editorial staff))

---

**Figure S1:**
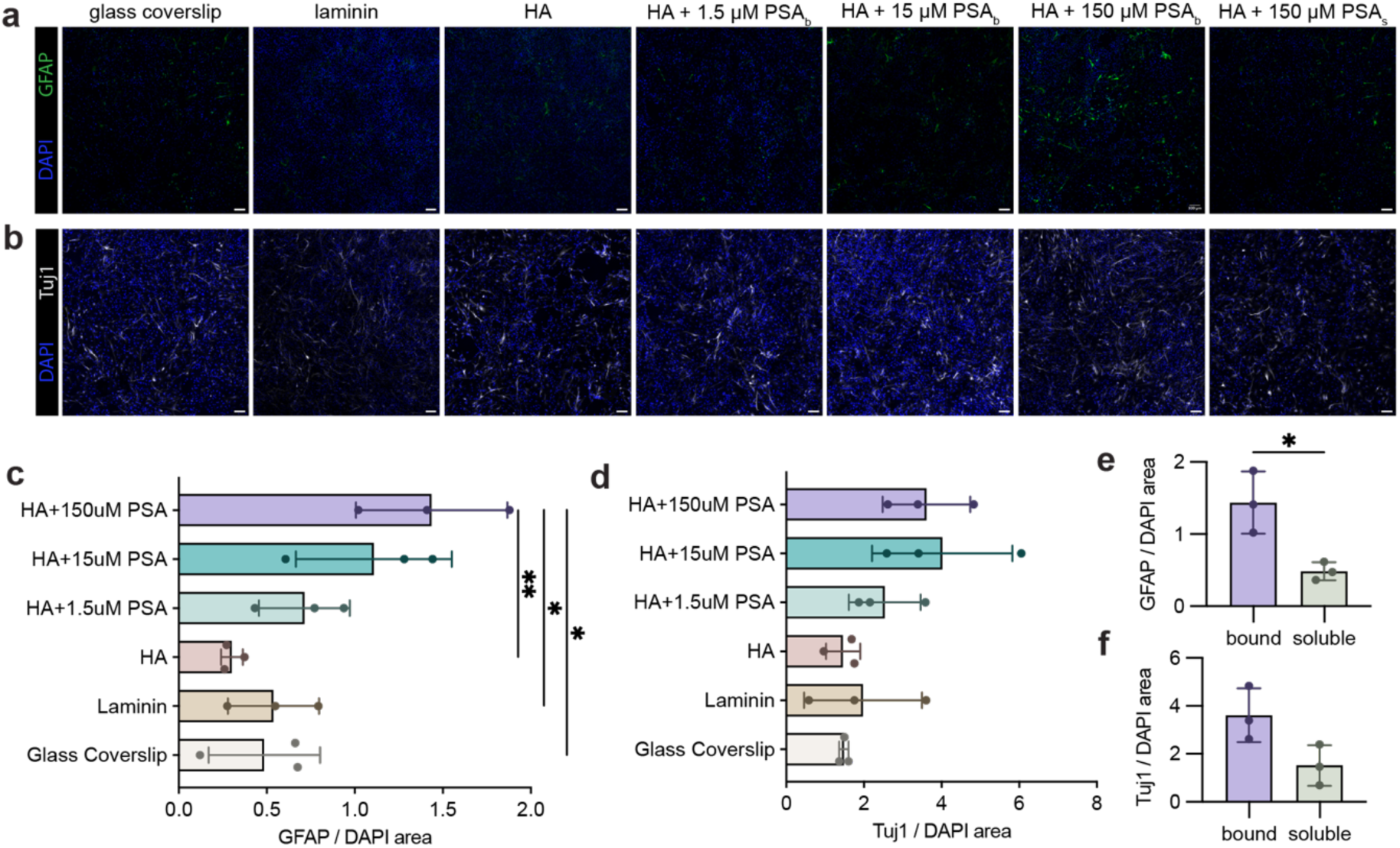
PSA on 2D coverslip biases NPC lineage markers in a dose- and presentation-dependent manner. a–b) Representative images of NPC stained with (a) GFAP with DAPI and (b) Tuj1 with DAPI. c–d) Quantification of (c) GFAP/DAPI area and (d) Tuj1/DAPI area across bound PSA conditions. e–f) Comparison of (e) GFAP/DAPI area and (f) Tuj1/DAPI area between bound and soluble PSA at 150uM concentration. Data are mean ± SEM. Each point is a biological replicate performed on a separate day from an independent cell vial; each biological replicate comprises two coverslips (technical replicates). Statistical tests: (c–d) one-way ANOVA with Tukey’s post-hoc test; (e–f) unpaired two-tailed t-test. *p < 0.05, **p < 0.01.

**Figure S2.**
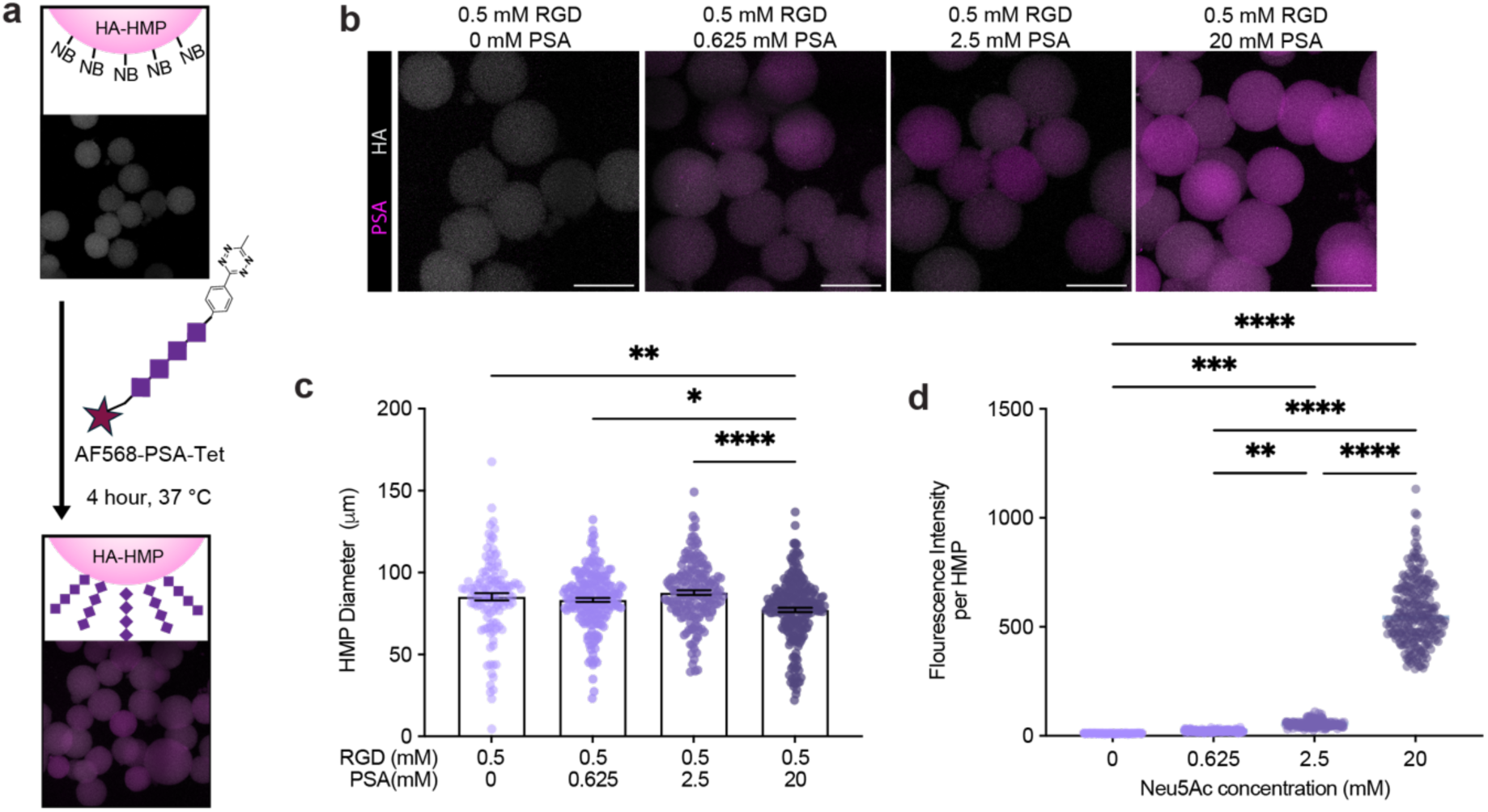
Characterization of PSA conjugation and microparticle size distributions. a) Schematic illustrating PSA conjugation onto HA-HMPs via tetrazine–norbornene chemistry, and representative fluorescent images confirming successful PSA conjugation at varying PSA concentrations. b) Representative fluorescent microscopy images of HMPs modified with 0 mM, 0.625 mM, 2.5 mM, and 20 mM PSA, scale bar: 100 µm. c) Quantitative measurements of microparticle diameters for the four PSA concentrations. d) Quantitative fluorescence intensity analysis demonstrates PSA conjugation densities on HMPs at increasing PSA concentrations. Data represent mean ± SEM, where each data point represents individual HMP. Statistical test: one-way ANOVA with Tukey’s post-hoc test, *p<0.05, **p<0.01, ****p<0.0001.

**Figure S3.**
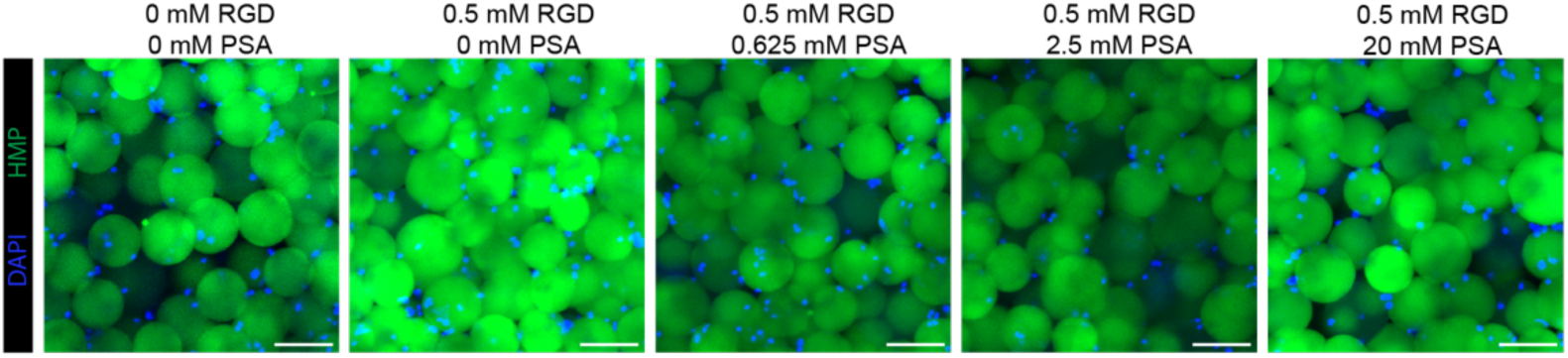
Representative confocal images of NPCs in scaffolds stained with DAPI on day 0.

**Figure S4:**
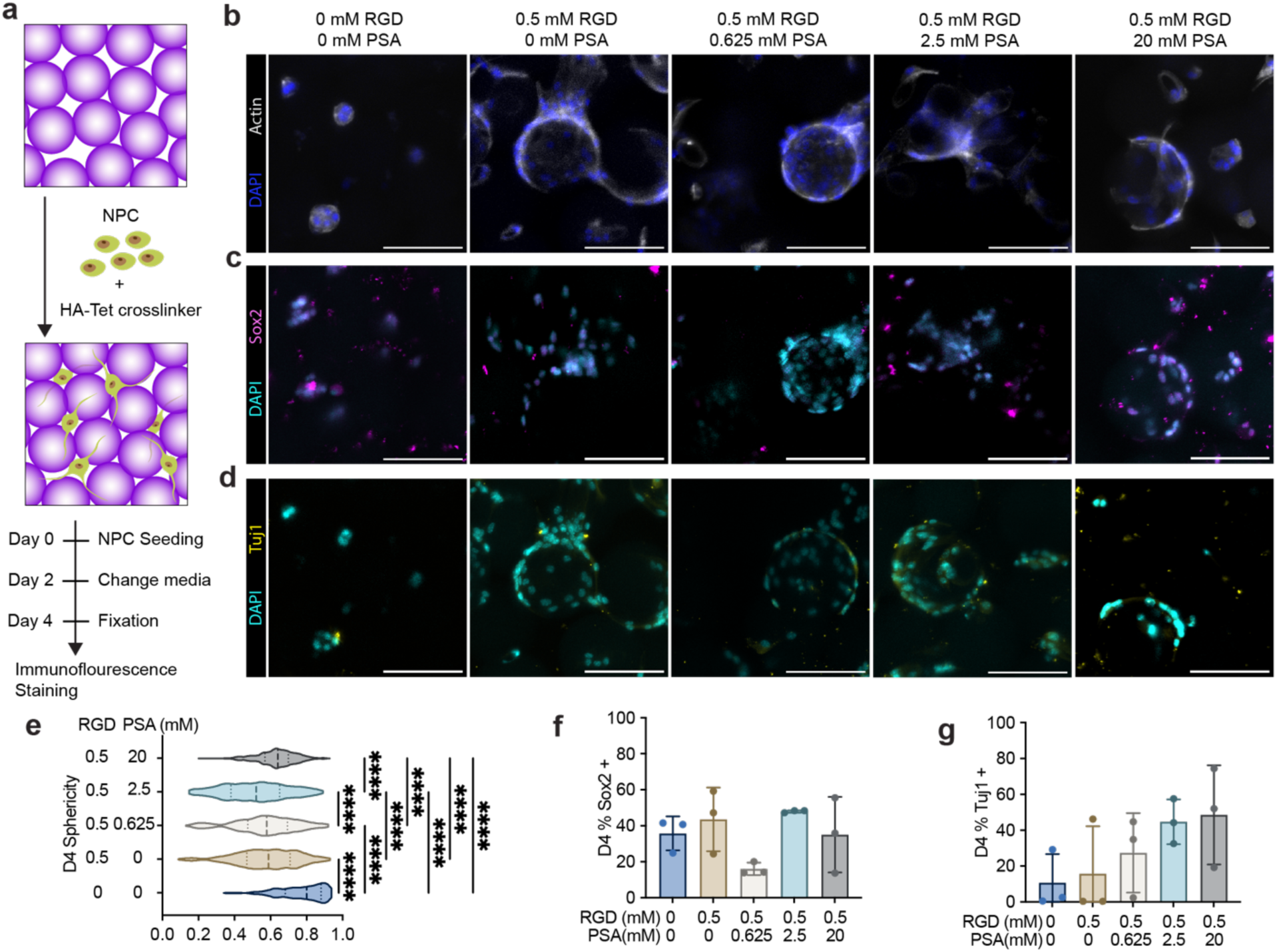
Effects of scaffold-tethered PSA on NPC morphology and progenitor marker expression at day 4. (a) Schematic of experimental setup illustrating NPC encapsulation within MAP scaffolds for a 4-day culture period. (b–d) Representative confocal images of NPCs in scaffolds stained with DAPI and actin (b), the progenitor marker SOX2 (c), or early neuronal differentiation marker Tuj1 (d). Scale bars: 100 µm. (e) Violin plots quantifying cell aggregate sphericity across scaffold conditions. Each data point represents an individual aggregate (>200 aggregates per condition, pooled from three biological replicates). (f,g) Quantification of the percentage of SOX2-positive (f) and Tuj1-positive (g) nuclei. Data represent mean ± SEM from three biological replicates,where each replicate corresponds to NPCs thawed from separate vials, scaffolds prepared independently, and experiments conducted on separate days (n = 3 scaffolds per condition). Statistical significance determined by one-way ANOVA with Tukey’s post-hoc test (*p < 0.05, **p < 0.01, ***p < 0.001, ****p < 0.0001).

**Figure S5.**
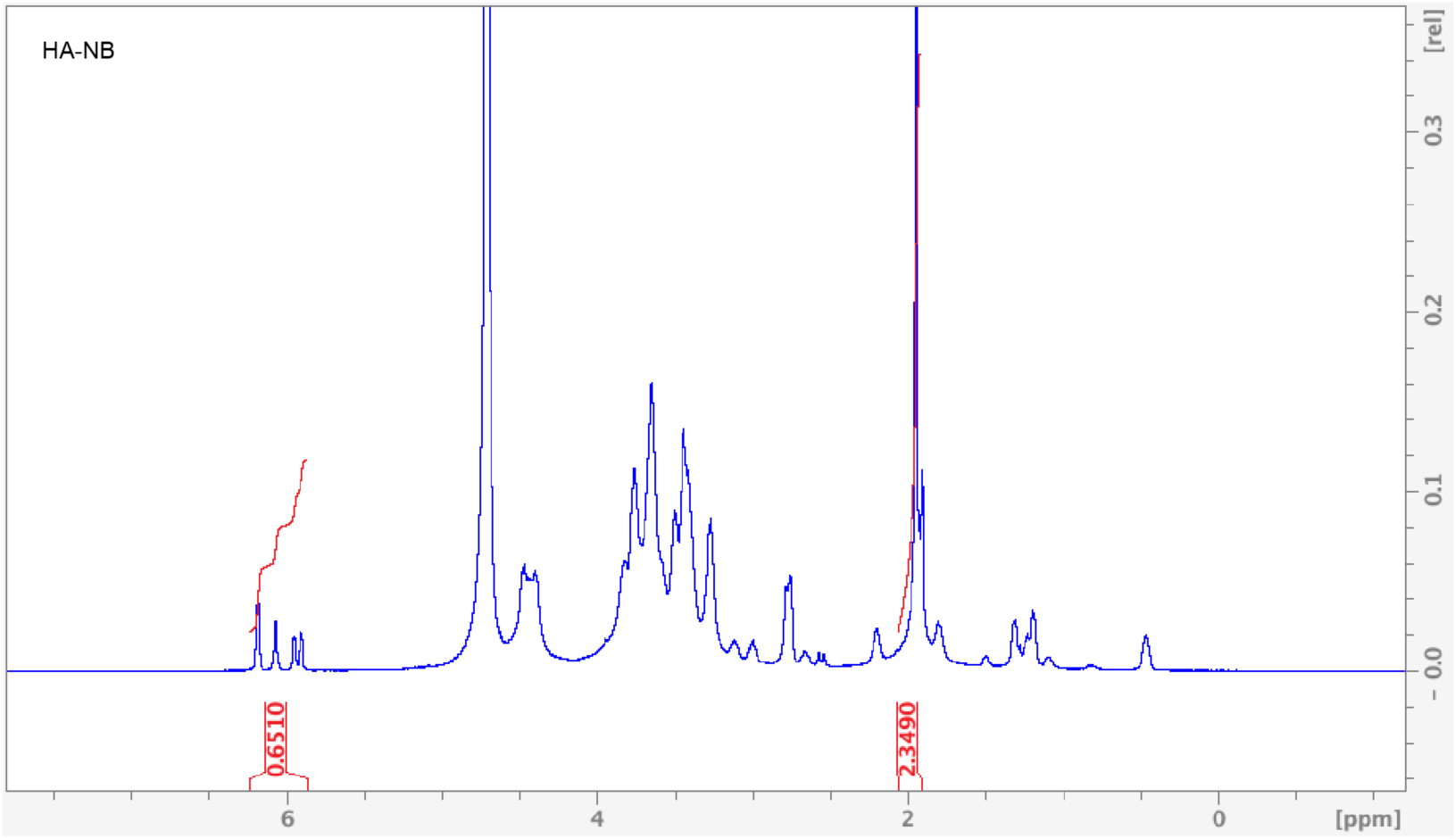
^1^H NMR analysis of HA-NB. ^1^H NMR signals in δ6.33 and δ6.02 (vinyl protons, endo), and δ6.26 and δ6.23 ppm (vinyl protons, exo) represent pendant norbornenes in D_2_O.

**Figure S6.**
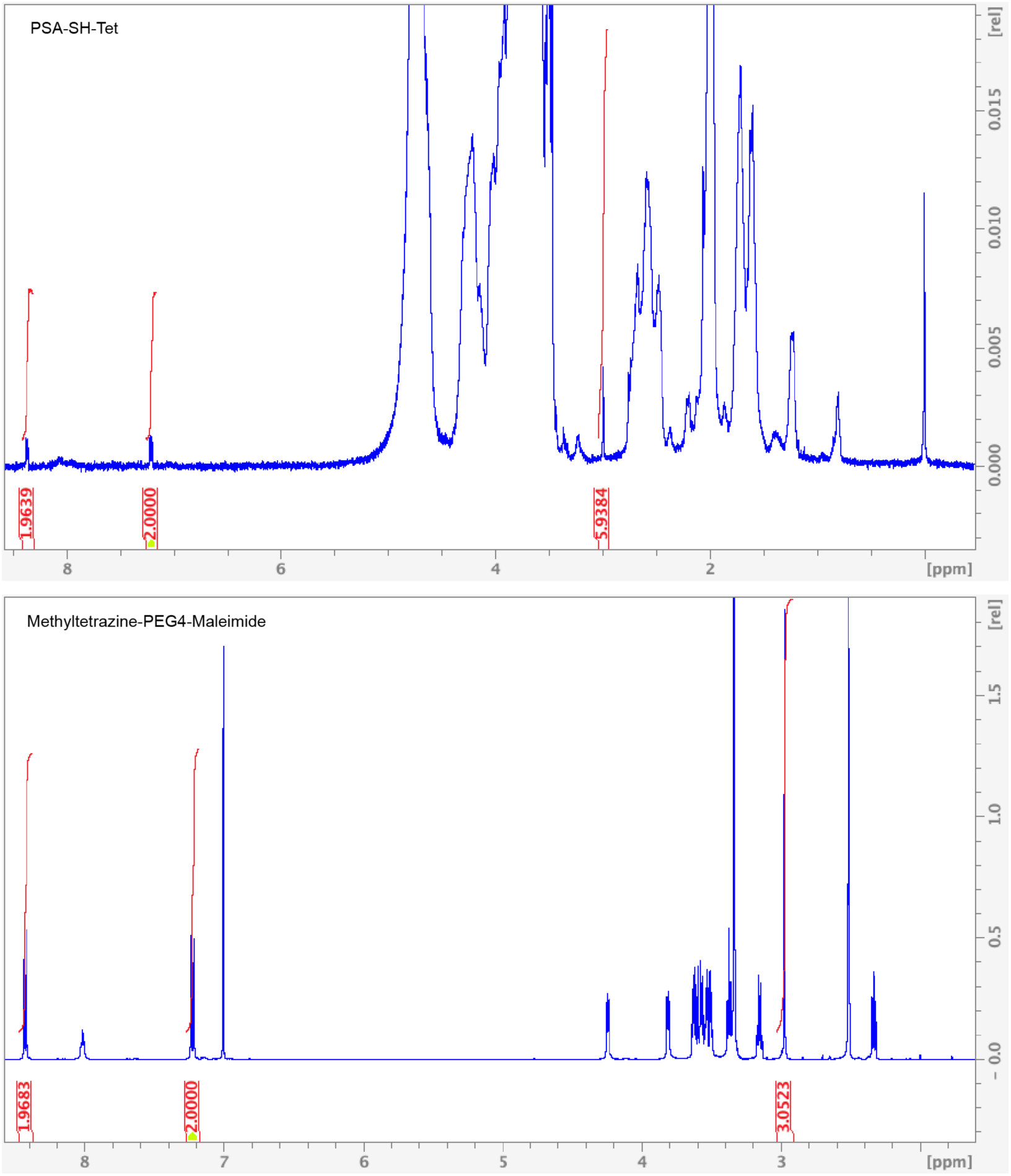
^1^H NMR analysis of PSA-Tet and methyltetrazine-PEG4-maleimide. ^1^H NMR signals at δ8.35 (2H) and δ7.20 (2H) (aromatic protons) represent tetrazines in D_2_O.

**Figure S7.**
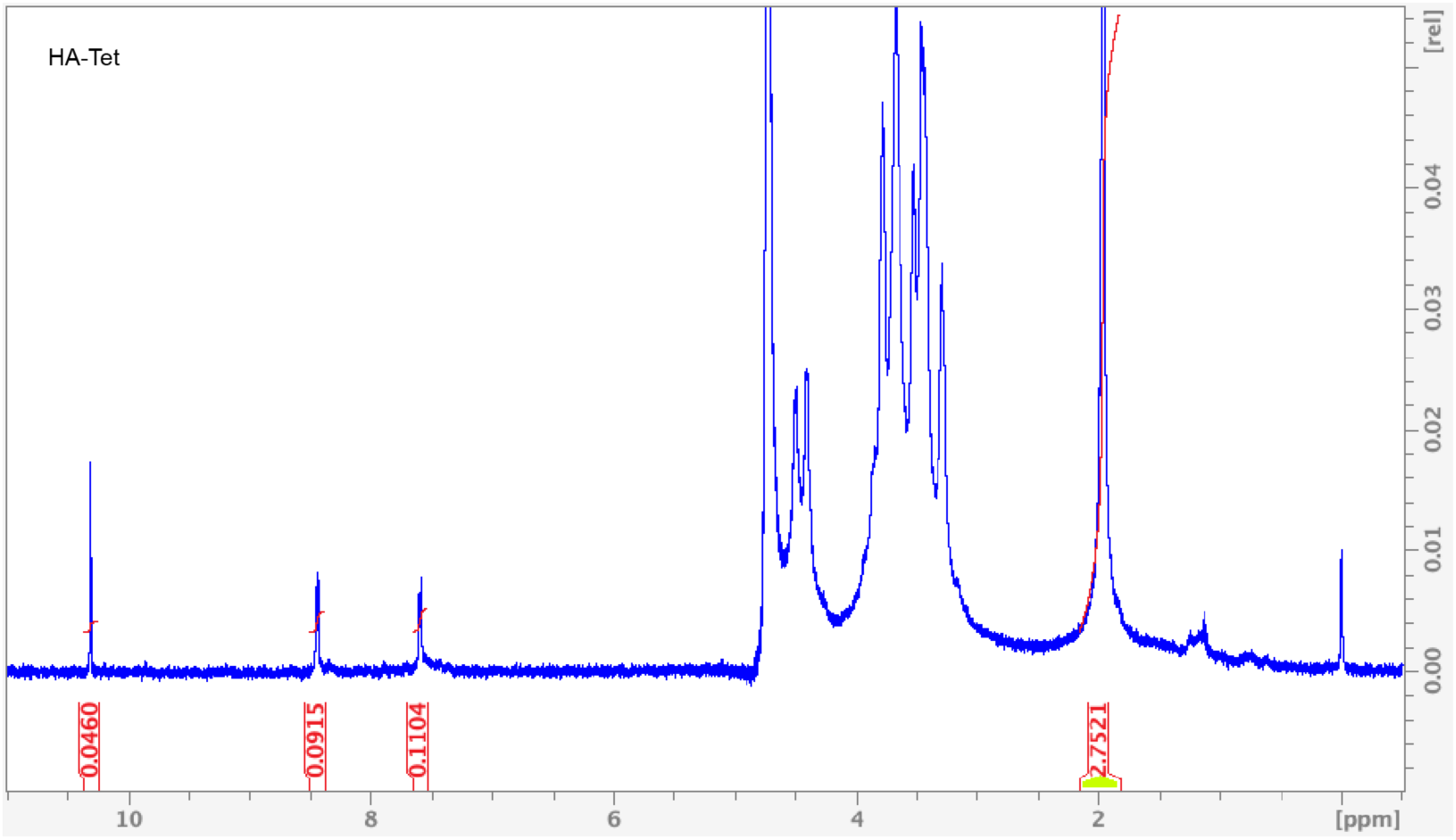
^1^H NMR analysis of HA-Tet. ^1^H NMR signals at δ8.5 (2H) and δ7.7 (2H) (aromatic protons) represent tetrazines in D_2_O.

**Figure S8.**
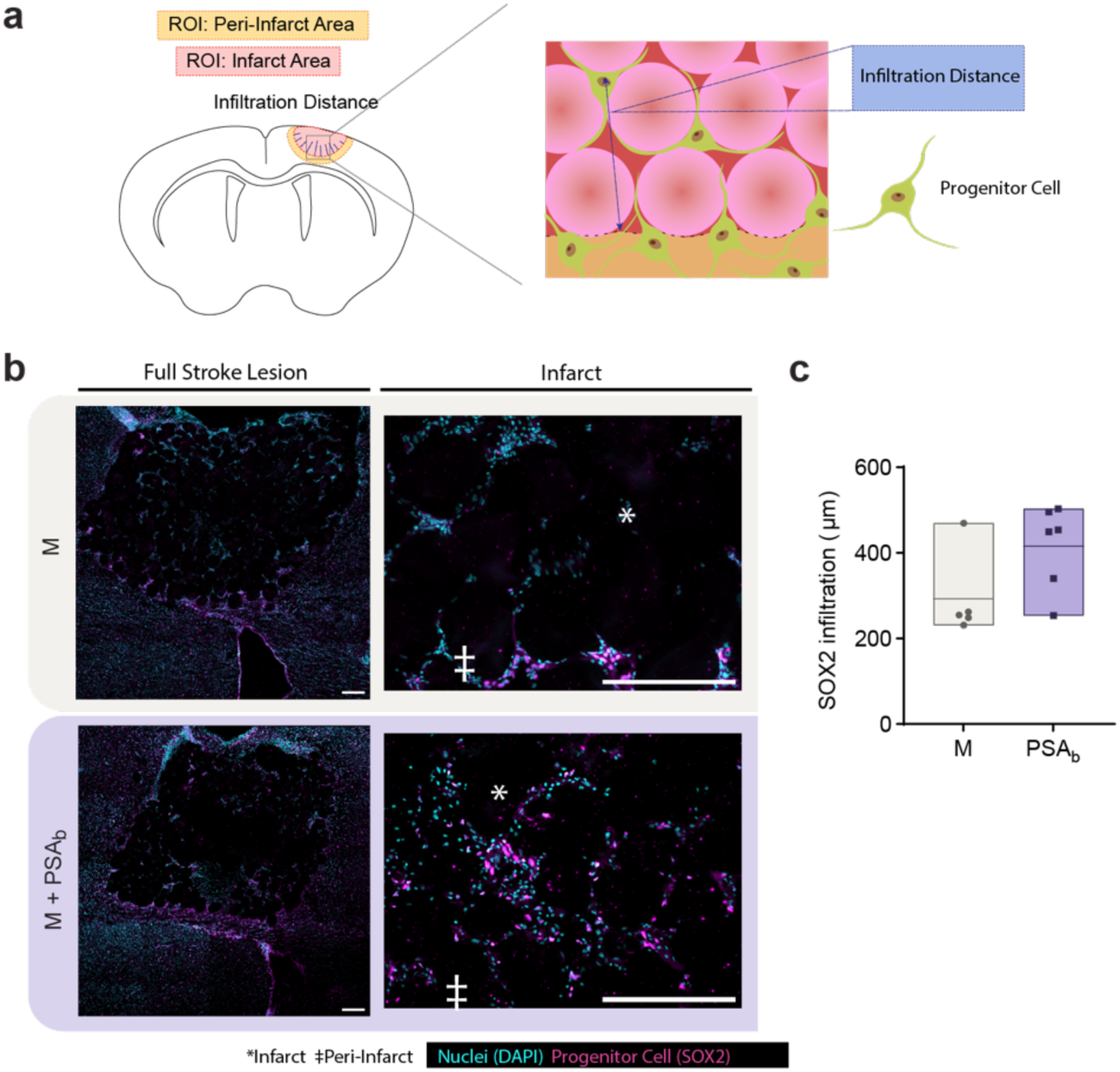
Quantification of Sox2 infiltration from peri-infarct border towards the infarct. a) Image analysis ROIs and schematic of infiltration distance measurement. b) Representative images of Sox2 staining for general endothelial cell visualization. c) Quantification of selected Sox2 signal distance the infarct border. Each data point is a biological replicate averaged from two coronal sections and plotted in a floating bar (n = 6). Two-tailed, unpaired Student’s t-test was performed. Scale bars represent 200 µm.

**Figure S9.**
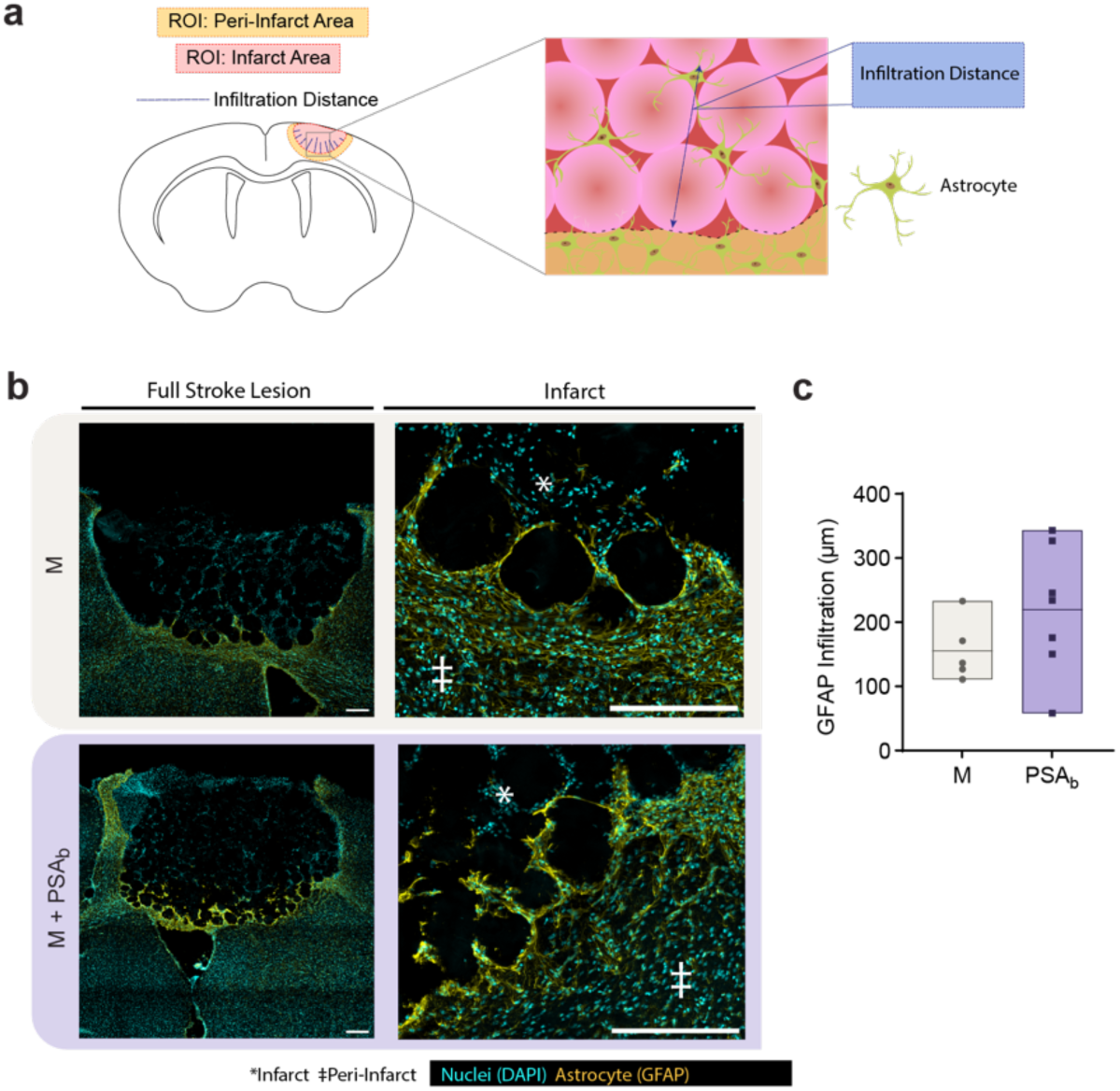
Quantification of GFAP infiltration from peri-infarct border towards the infarct. a) Image analysis ROIs and schematic of infiltration distance measurement. b) Representative images of GFAP staining for general endothelial cell visualization. c) Quantification of selected GFAP signal distance the infarct border. Each data point is a biological replicate averaged from two coronal sections and plotted in a floating bar (n = 6). Two-tailed, unpaired Student’s t-test was performed. Scale bars represent 200 µm.

**Figure S10.**
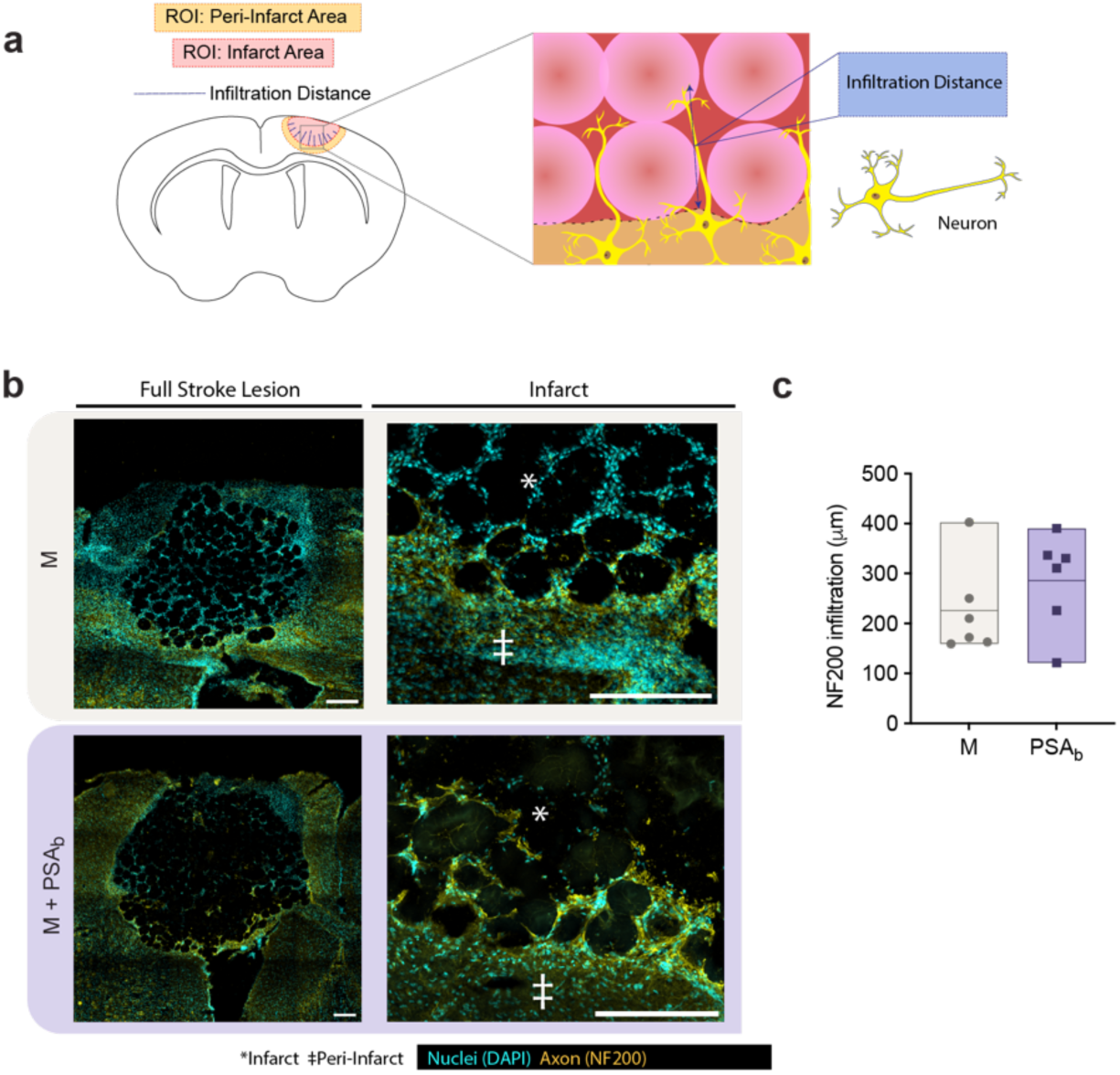
Quantification of NF200 infiltration from peri-infarct border towards the infarct. a) Image analysis ROIs and schematic of infiltration distance measurement. b) Representative images of NF200 staining for general endothelial cell visualization. c) Quantification of selected NF200 signal distance the infarct border. Each data point is a biological replicate averaged from two coronal sections and plotted in a floating bar (n = 6). Two-tailed, unpaired Student’s t-test was performed. Scale bars represent 200 µm.

**Figure S11.**
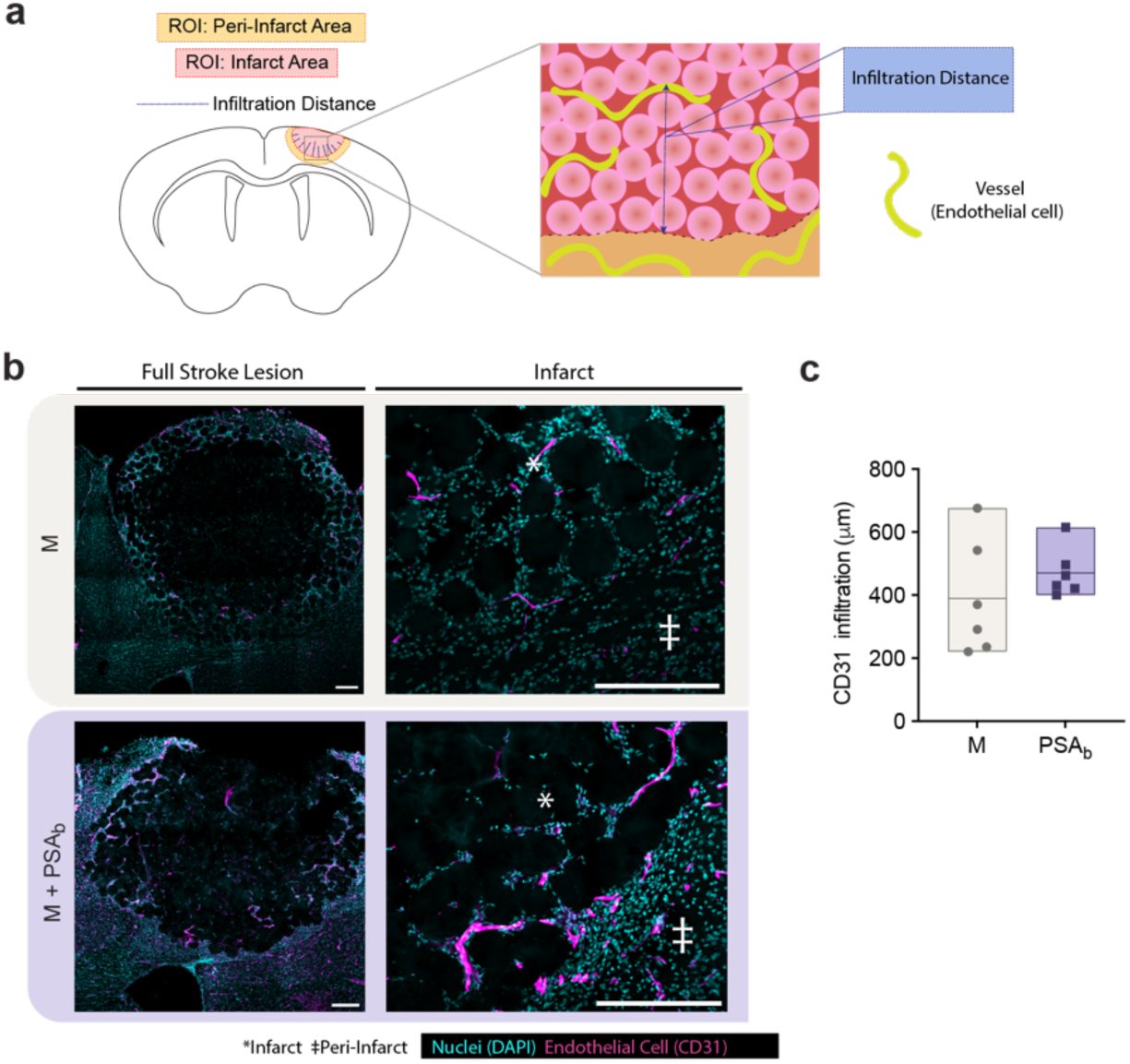
Quantification of CD31 infiltration from peri-infarct border towards the infarct. a) Image analysis ROIs and schematic of infiltration distance measurement. b) Representative images of CD31 staining for general endothelial cell visualization. c) Quantification of selected CD31 signal distance the infarct border. Each data point is a biological replicate averaged from two coronal sections and plotted in a floating bar (n = 6). Two-tailed, unpaired Student’s t-test was performed. Scale bars represent 200 µm.

